# Developmental constraints on genome evolution in four bilaterian model species

**DOI:** 10.1101/161679

**Authors:** Jialin Liu, Marc Robinson-Rechavi

**Affiliations:** Department of Ecology and Evolution, University of Lausanne, Switzerland; Swiss Institute of Bioinformatics, Lausanne 1015, Switzerland

## Abstract

Developmental constraints on genome evolution have been suggested to follow either an early conservation model or an “hourglass” model. Both models agree that late development strongly diverges between species, but debate on which developmental period is the most conserved. Here, based on a modified “Transcriptome Age Index” approach, i.e. weighting trait measures by expression level, we analyzed the constraints acting on three evolutionary traits of protein coding genes (strength of purifying selection on protein sequences, phyletic age, and duplicability) in four species: nematode worm *Caenorhabditis elegans*, fly *Drosophila melanogaster*, zebrafish *Danio rerio*, and mouse *Mus musculus*. In general, we found that both models can be supported by different genomic properties. Sequence evolution follows an hourglass model, but the evolution of phyletic age and of duplicability follow an early conservation model. Further analyses indicate that stronger purifying selection on sequences in the middle development are driven by temporal pleiotropy of these genes. In addition, we report evidence that expression in late development is enriched with retrogenes, which usually lack efficient regulatory elements. This implies that expression in late development could facilitate transcription of new genes, and provide opportunities for acquisition of function. Finally, in *C. elegans*, we suggest that dosage imbalance could be one of the main factors that cause depleted expression of high duplicability genes in early development.

## Introduction

Evolutionary changes in the genome can cause changes in development, which are subject to natural selection. This leads developmental processes to constrain genome evolution. More precisely, selection on the output of development affects evolution of the genomic elements active in development. Currently, based on morphological similarities during development, two popular models have been proposed to bridge developmental and evolutionary biology. The early conservation model, modified from the “third law” of Von Baer (1828) (as cited in Kalinka and Tomancak 2012), suggests that the highest morphological similarities among species from the same phylum occur in early development, followed by a progressive evolutionary divergence over ontogeny. It should be noted that Von Baer in fact based his observations on post-gastrulation embryos (Kalinka and Tomancak 2012; Abzhanov 2013). The “developmental burden” concept was proposed to explain this model. It suggested that the development of later stages is dependent on earlier stages, so that higher conservation should be found in the earlier stages of development (Garstang 1922; Riedl 1978) (as discussed in Irie and Kuratani 2014). Based on renewed observations in modern times, however, Duboule (1994) and Raff (1996) proposed the developmental “hourglass model”. This model suggested that a “phylotypic period” (Richardson 1995) in middle development has higher morphological similarities than early or late development. Several mechanisms have been proposed to explain this observation. Duboule (1994) proposed that it may be due to co-linear Hox cluster gene expression in time and space. Raff (1996) suggested a high inter-dependence in signaling among developmental modules in middle development. Galis and Metz (2001) also highlighted the high number of interactions at this period, although Comte et al. (2010) did not find any molecular evidence for these interactions. It is worth noting that the hourglass model was not supported by a comprehensive study of vertebrate embryonic morphology variation (Bininda-Emonds et al. 2003). A number of alternatives have been proposed, for example the “adaptive penetrance model” (Richardson et al. 1997) and the “ontogenetic adjacency model” (Poe and Wake 2004). Of note, the higher divergence of late development could also be due to stronger adaptive selection, the “Darwin hypothesis” (Artieri et al. 2009). This question is independent of the pattern of constraints (early or hourglass) which are the focus of this study, and we explore it in a companion study (Liu and Robinson-Rechavi 2017).

Both main models have been supported by recent genomic level studies based on different properties (such as expression divergence, sequence divergence, duplication, or phyletic age), different species, and different analysis methods. Concerning expression divergence, interestingly, all studies are consistent across different species and research groups (Kalinka et al. 2010; Irie and Kuratani 2011; Yanai et al. 2011; Levin et al. 2012; Wang et al. 2013; Gerstein et al. 2014; Ninova et al. 2014; Zalts and Yanai 2017). All of them suggested that middle development has the highest transcriptome conservation, i.e. the hourglass pattern. On the other hand, when animals are compared between different phyla, middle development has been reported to have the highest divergence (Levin et al. 2016), although this conclusion has been criticized on methodological grounds (Dunn et al. 2018). From other properties, however, the results are inconclusive based on different methods (Castillo-Davis and Hartl 2002; Cutter and Ward 2005; Davis et al. 2005; Hazkani-Covo et al. 2005; Hanada et al. 2007; Irie and Sehara-Fujisawa 2007; Cruickshank and Wade 2008; Roux and Robinson-Rechavi 2008; Artieri et al. 2009; Domazet-Loso and Tautz 2010; Quint et al. 2012; Piasecka et al. 2013; Cheng et al. 2015; Drost et al. 2015).

Generally, the methods used to measure developmental constraints at the genomic level can be divided into three categories: proportion based analysis, module analysis, and transcriptome index analysis.

Proportion based analysis consists in testing the proportion of genes with a given property within all expressed genes (Roux and Robinson-Rechavi 2008). The method is less used following the emergence of accurate transcriptome-scale data, since it does not take into account the contributions of expression abundance.

Module analysis consists in studying evolutionary properties of distinct sets of genes (modules) which are specifically expressed in groups of developmental stages (Piasecka et al. 2013). This method can avoid problems caused by genes expressed over all or a large part of development. For example, trends might be diluted by highly expressed housekeeping genes, which contribute to the average expression at all developmental stages. However, this approach can only measure the developmental constraints for a specific subset of genes, instead of considering the composition of the whole transcriptome.

Transcriptome index analysis is a weighted mean: the mean value of an evolutionary parameter is weighted by each gene’s expression level (Domazet-Loso and Tautz 2010). This method has the benefit of detecting evolutionary constraints on the whole transcriptome, but patterns can be driven by a subset of very highly expressed genes, or even by a few outliers, because the difference between highly and lowly expressed genes can span several orders of magnitude. For instance, Domazet-Loso and Tautz (2010) reported that transcriptomes of middle development stages of *D. rerio* have a higher proportion of old genes than transcriptomes of early and late development stages, using the transcriptome age index. However, Piasecka et al. (2013) re-analyzed the same data and reported that the highest proportion of old genes was in transcriptomes of early development stages, once a standard log-transformation of microarray signal intensities was done, a result confirmed by module analysis and proportion based analysis.

In addition, several statistical methods have been proposed to distinguish the hourglass model from the early conservation model.

The parabolic test is based on fitting both first degree and second degree polynomial models (Roux and Robinson-Rechavi 2008). The hourglass model is supported if the parabolic function provides a significantly better fit and its minimum corresponds to middle development. This method has been criticized for being too specific and insensitive to other non-parabolic hourglass patterns (Drost et al. 2015).

The flat line test simply tests whether variance of transcriptome indexes across development is significantly higher than variance from random samples (Domazet-Loso and Tautz 2010; Quint et al. 2012). But a significant difference does not necessarily imply the existence of an hourglass pattern (Drost et al. 2015).

Since these two methods are either too strict or without power to distinguish the hourglass model, Drost et al. (2015) proposed a “reductive hourglass test” which focuses on testing the presence of an hourglass pattern of divergence: high-low-high. For this, development can be divided into three periods (early, phylotypic, and late), based on the known phylotypic period from morphological studies. Then, a permutation method is used to test whether the mean value in the phylotypic period is significantly lower than in early and late periods.

Overall, the transcriptome index analysis should be the best method to measure developmental constraints on the whole transcriptome, if care is taken to properly treat the expression values. Moreover, the reductive hourglass test should be used to objectively test the hourglass model, alone or in combination with other methods.

Because previous studies used different methodologies, and few studies adopted a transformed transcriptome index analysis, their conclusions cannot be compared consistently, making a biological conclusion concerning developmental constraints across species and features difficult. What’s more, while many studies focus on distinguishing between early conservation model and hourglass conservation model, we still know very little of the factors driving these patterns.

To measure developmental constraints on genome evolution, we calculated transcriptome indexes over the development of four species (*C. elegans, D. melanogaster, D. rerio and M. musculus*), for three evolutionary parameters (strength of purifying selection on coding sequences (ω_0_), phyletic age, and duplicability (paralog number)), with three transformations of expression values (non-transformed, log_2_ transformed, and square root transformed). For *C. elegans*, the strength of purifying selection on coding sequences was not reliably estimated, with no data in the Selectome database (Moretti et al. 2014) and very high values of estimated synonymous distances (dS) from Ensembl Metazoa (Kersey et al. 2016) (Figure S1); thus we did not include this parameter in the study of *C. elegans*. In general, we found results consistent with the hourglass model for sequence evolution, but with early conservation for phyletic age and paralog number, in the four species. In addition, log_2_ transformed transcriptome indexes are always consistent with square root transformed transcriptome indexes but not with non-transformed transcriptome indexes.

## Results and discussion

### Effect of expression value transformation on transcriptome indexes

As mentioned in the Introduction, the pattern from a transcriptome index analysis may not reflect the global behavior of the transcriptome, but that of a small fraction of very highly expressed genes, or even of a few outliers. In order to systematically test this issue, we calculated 95% confidence intervals of transcriptome indexes based on log_2_ transformed, square root transformed, and non-transformed expression values (see Methods). Then, for the purpose of comparing the range of confidence intervals in the same scale, we plotted the ratio of upper to lower confidence interval boundary across development. Clearly, at a given confidence level (95% here), we can see that the ratio of non-transformed transcriptome indexes is much higher and more variable than transformed transcriptome indexes (Figure S2), indicating that the transcriptome indexes estimated from transformed expression are more stable. The most stable pattern comes from log_2_ transformed transcriptome indexes, although it is quite similar with square root transformation. We note that while log-transformation is routine in most application of transcriptomics, many analyses of “hourglass” patterns use non transformed expression data.

In summary, although a subset of genes with dramatically different expression values in different stages could be interesting in some sense, when the goal is to investigate the general tendency of the transcriptome, log-or square-root-transformation for expression value is necessary and efficient to reach a stable estimation.

### Variation of evolutionary transcriptome indexes across development

Here, based on log_2_ transformed expression values, we calculated transcriptome indexes for strength of purifying selection on coding sequences (ω_0_), phyletic age, and duplicability (paralog number). In order to objectively distinguish the hourglass model from the early conservation model, we used a permutation test method similar to that of Drost et al. (2015) (see Methods). For all parameters considered the highest divergence is observed in late development, and there are many more stages sampled from late development, so we only compared the difference between early and middle development. Thus a significant *p-*value for lower divergence in middle *vs*. early development supports the hourglass model, whereas a lack of significance supports the early conservation model. We consider early conservation to cover both stronger conservation in early than middle development, and similar strong conservation over early and middle development, and hence we use a one-sided test. Notably, for early development, we did not consider the stages before the start of maternal to zygote transition (MZT, see Methods), because these stages are dominated by maternal transcripts.

For the transcriptome index of purifying selection on coding sequence (Transcriptome Divergence Index: TDI), we found that genes with stronger purifying selection tend to be more expressed at middle developmental stages, suggesting an hourglass pattern (Figure 1). However, for the transcriptome indexes of phyletic age (Transcriptome Age Index: TAI) and of paralog number (Transcriptome Paralog Index: TPI), we observed that genes with higher duplicability and younger phyletic age trend to be expressed at later developmental stages, which corresponds to early conservation (Figure 1). In addition, we also repeated these analyses based on square root transformed expression values (Figure S3) and on non-transformed expression values (Figure S4). In general, the results from square root transformation are highly consistent with those from log_2_ transformation, but not with those from non-transformation. For example, with non-transformed expression data, the TPI in *C. elegans* became very noisy; the TAI in *D. rerio* changed from early conservation to hourglass pattern; or the TDI in *M. musculus* changed into an unexpected early divergence pattern. Finally, we confirmed our observations with other datasets (Figure S5). For *C. elegans, D. melanogaster* and *D. rerio*, we used a high resolution time series single embryo RNA-seq dataset (Levin et al. 2016). Since this dataset is without replicates, and is generated from single embryo, the transcriptome indexes are noisy and present extreme values in some time points. However, generally, all the results from the new dataset are consistent with the results from our previous datasets except the TDI in *D. rerio.* For the latter, we only observed the first two-thirds of an hourglass pattern in the new dataset. This is because the new dataset only covers embryo development, whereas the increased TDI in late development is driven by post-embryonic development stages. For *M. musculus*, we also confirmed our results based on a microarray dataset (Irie and Kuratani 2011) (Figure S5).

**Figure 1:**
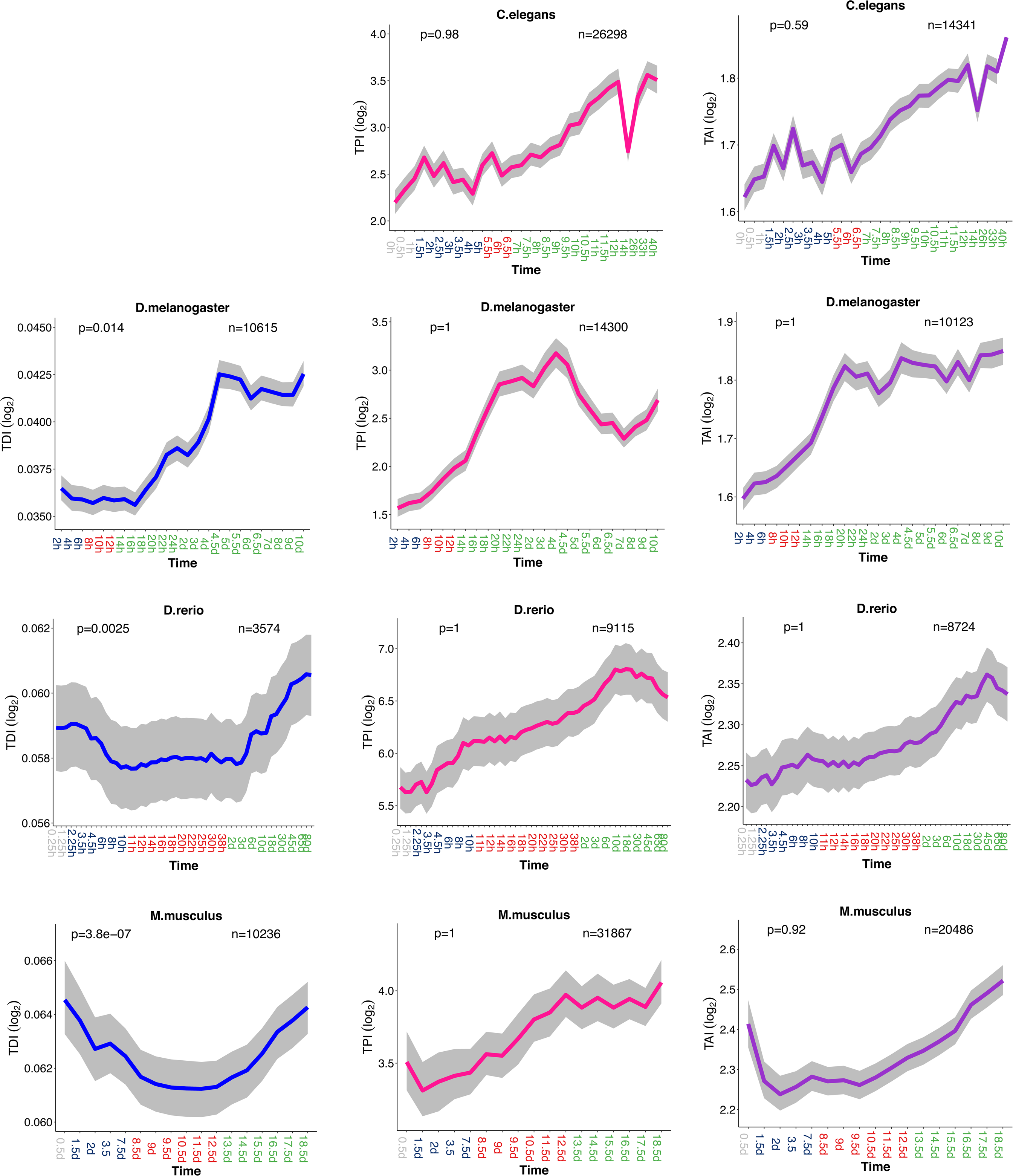
Evolutionary transcriptome indexes based on log_2_ transformed expression values. Grey, dark blue, red, and green marked time points in the x-axis represent stages before the start of MZT, early developmental stages, middle developmental stages, and late developmental stages, respectively. Transcriptome index of divergence (TDI): blue line; transcriptome index of paralog number (TPI): pink line; transcriptome index of phyletic age (TAI): purple line. The grey area indicates 95% confidence interval estimated from bootstrap analysis. The *p-*values for supporting the hourglass model (permutation test, early *vs*. middle development) are indicated in the top-left corner of each plot. The numbers of genes analyzed are noted in the top-right corner of each plot.

In *D. melanogaster*, we did not confirm the results of Drost et al. (2015) for phyletic age. After log_2_ transformation of expression data, we found an early conservation pattern instead of the hourglass pattern which they reported (Figure S6B). It appears that the hourglass pattern of phyletic age in their study is driven by a few highly expressed genes, consistently with our previous observations in *D. rerio* (Piasecka et al. 2013). This is verified by excluding the top 10% most expressed genes and analyzing without transformation (Figure S6C). Of note, that fly phyletic age hourglass with uncorrected expression was also reported earlier (Domazet-Loso and Tautz 2010).

Overall, these results suggest that genes under strong purifying selection on their protein sequence trend to be expressed in middle development; it remains to be seen how much these observations extend to more arthropods or chordates. They also extend our previous observations that genes expressed earlier have a lower duplicability and an older age (Roux and Robinson-Rechavi 2008; Piasecka et al. 2013). In addition, it poses the question whether a pattern driven by the minority of very highly expressed genes is relevant to understanding Evo-Devo, which is generally driven by regulatory genes (Carroll 2008), such as transcription factors, with typically not very high and rather tissue-specific expression.

### Expression of temporal pleiotropy genes across development

Several models have been proposed to explain why some developmental stages are more conserved than others, as presented in the Introduction. In all models, a common point is that high conservation is caused by selection against deleterious pleiotropic effects of mutations. This implies that higher sequence conservation in middle developmental stages is caused by higher pleiotropy of genes expressed in these stages, pleiotropy being one of the major factors that constrain sequence evolution (Fraser et al. 2002).

In order to test this hypothesis, we used one type of development related pleiotropic effect: temporal pleiotropy (Artieri et al. 2009) (expression breadth across development). This is similar to spatial pleiotropy (Larracuente et al. 2008; Kryuchkova-Mostacci and Robinson-Rechavi 2015) (expression breadth across tissues) or connective pleiotropy (Fraser et al. 2002) (protein-protein connectivity). The more stages a gene is expressed in, the more traits it could affect, so it is expected to be under stronger evolutionary constraints (Wagner and Zhang 2011). For *C. elegans, D. melanogaster and M. musculus*, we defined FPKM>1 as expressed. For *D. rerio* we set genes with microarray signal rank in top 70% as expressed.

We calculated the proportion of potentially pleiotropic genes as expressed in more than 50% of development stages. In all the species, interestingly, we found pleiotropic genes enriched in middle development (Figure 2). In *D. melanogaster*, the evidence is weaker, because of the low sampling of early and middle development stages in the main dataset; but the pattern was clear in the high resolution single embryo RNA-seq dataset (Figure S7). We also found similar patterns when we define pleiotropic genes as expressed in more than 70% of development stages (Figure S8). Since the late development of *D. melanogaster* can clearly be divided into two periods with distinct patterns from the pleiotropy analysis, we removed the second period of late development, and found the same overall trend (Figure S9). For *D. rerio*, in addition, we observed consistent results based on setting expressed genes as microarray signal rank in the top 90% or 50% (Figure S10). Similar observations of higher temporal pleiotropy for genes in middle development in vertebrates were recently reported by Hu et al. (2017).

**Figure 2:**
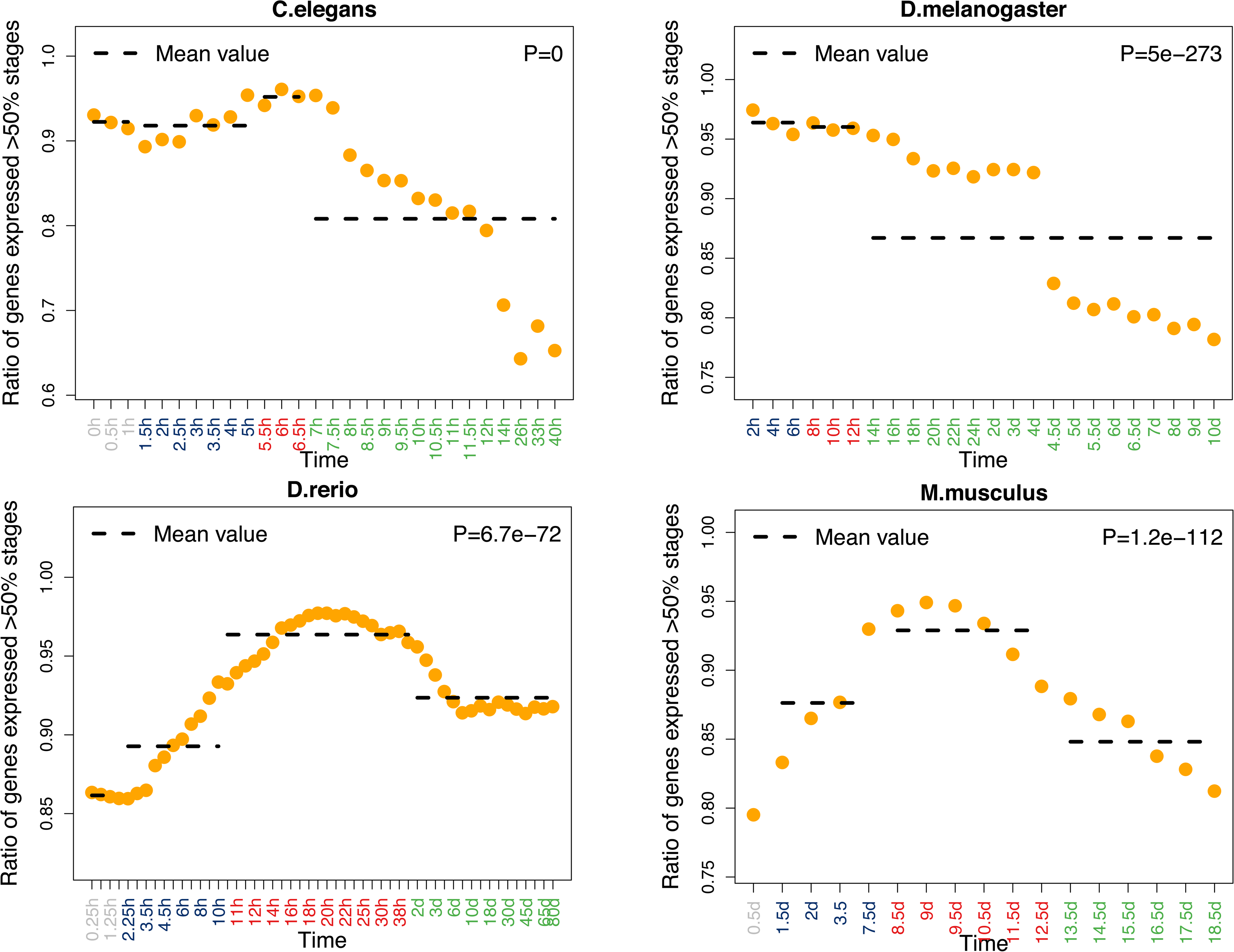
Proportion of temporal pleiotropic genes across development. Grey, dark blue, red, and green marked time points in the x-axis represent stages before the start of MZT, early developmental stages, middle developmental stages and late developmental stages respectively. The proportion of temporal pleiotropic genes is plotted as orange circles. The *p-*values from chi-square goodness of fit test are indicated in the top-right corner of each graph. Pleiotropic genes are defined as expressed in more than 50% of stages sampled. The proportion of pleiotropic genes is defined as the number of pleiotropic genes divided by the number of all genes expressed in the corresponding stage.

Based on these observations, we further checked whether higher temporal pleiotropic constraint could explain stronger purifying selection on sequence evolution. As expected, we found that pleiotropic genes have lower ω_0_ than non-pleiotropic genes (Figure 3).

**Figure 3:**
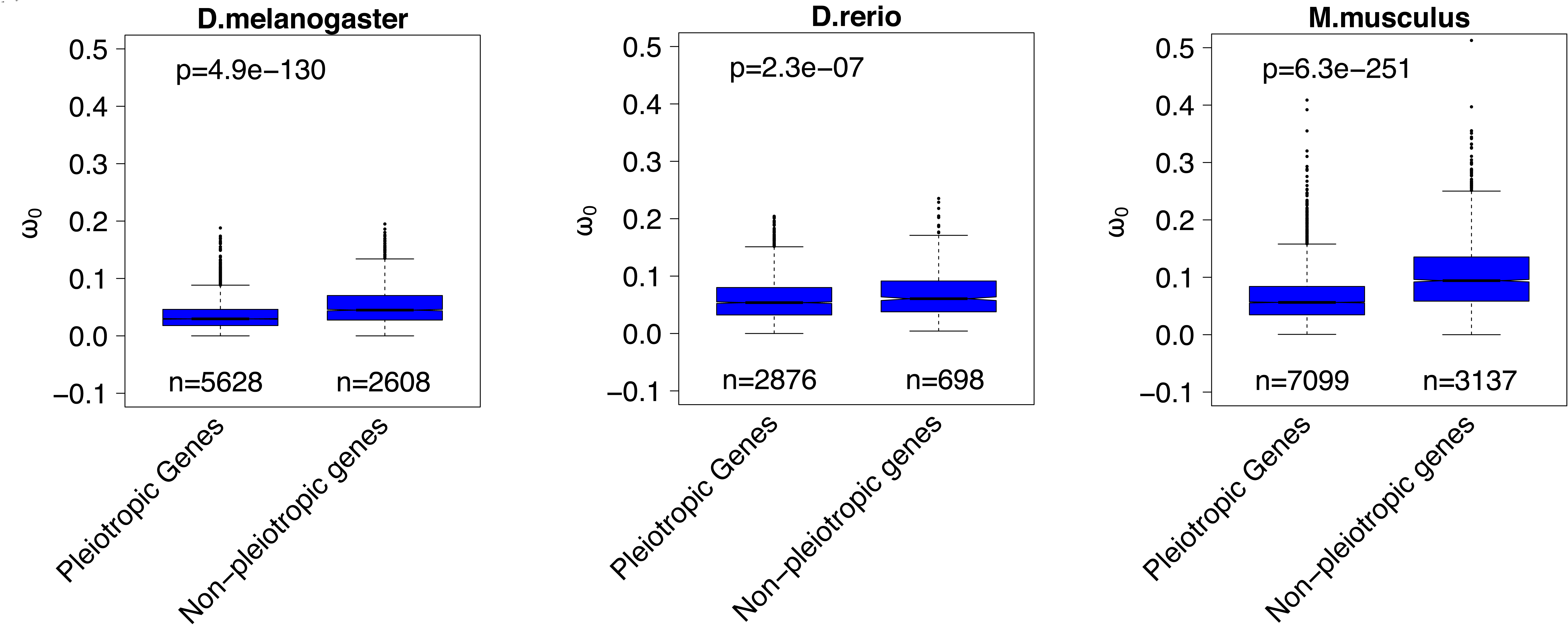
Comparison of ω_0_ between temporal pleiotropic genes and non-pleiotropic genes. The number of genes in each category is indicated below each box. The *p*-values from a Wilcoxon test comparing categories are reported above boxes. The lower and upper intervals indicated by the dashed lines (“whiskers”) represent 1.5 times the interquartile range, or the maximum (respectively minimum) if no points are beyond 1.5 IQR (default behaviour of the R function boxplot).

In summary, we found that middle development stages with a higher proportion of broadly expressed genes are under stronger pleiotropic constraint on sequence evolution.

### Higher expression of retrogenes in later development stages

In adult anatomy, young genes are mainly enriched for expression in testis (Kaessmann 2010). Two main factors have been proposed to explain this pattern. Firstly, permissive chromatin in testis facilitates the transcription of most genes, including new genes (Soumillon et al. 2013). The widespread expression in testis appears related to regulation of gene evolution rates based on transcription coupled repair (Xia et al. 2018). Secondly, as the most rapidly evolving organ at genomic level, there is least purifying selection acting on new genes expressed in testis (Kaessmann 2010). Is there a similar explanation for the ontogenic pattern of young genes tending to be expressed in late development stages? As testis constitutes the most rapidly evolving organ transcriptome, late development represents the most rapidly evolving stage transcriptome, owing to both relaxed purifying selection (Artieri et al. 2009) and to increased positive selection (Liu and Robinson-Rechavi 2017). Thus, we suggest that expression in late development might, like in testis, promote the fixation and functional evolution of new genes.

In order to test this, we analyzed the expression of retrogenes across development. Since retrogenes usually lack regulatory elements, most of them fail to acquire transcription and achieve function (Kaessmann et al. 2009). So, if late development, like testis, can facilitate the transcription of new genes, promoting their fixation, we should observe higher expression of retrogenes in later developmental stages. Because retrogenes have higher expression in testis, and testis is already differentiated after middle development, we excluded testis genes in our analyses for *D. melanogaster* and *M. musculus,* where the information of testis gene expression was available. As expected, the median expression of retrogenes is higher in late development (Figure 4), with a significant positive correlation. Generally, in *C. elegans, D. rerio and M. musculus*, the median expression progressively increases; in *D. melanogaster*, all the median values are 0 until stage 4 days, and then it progressively increases.

**Figure 4:**
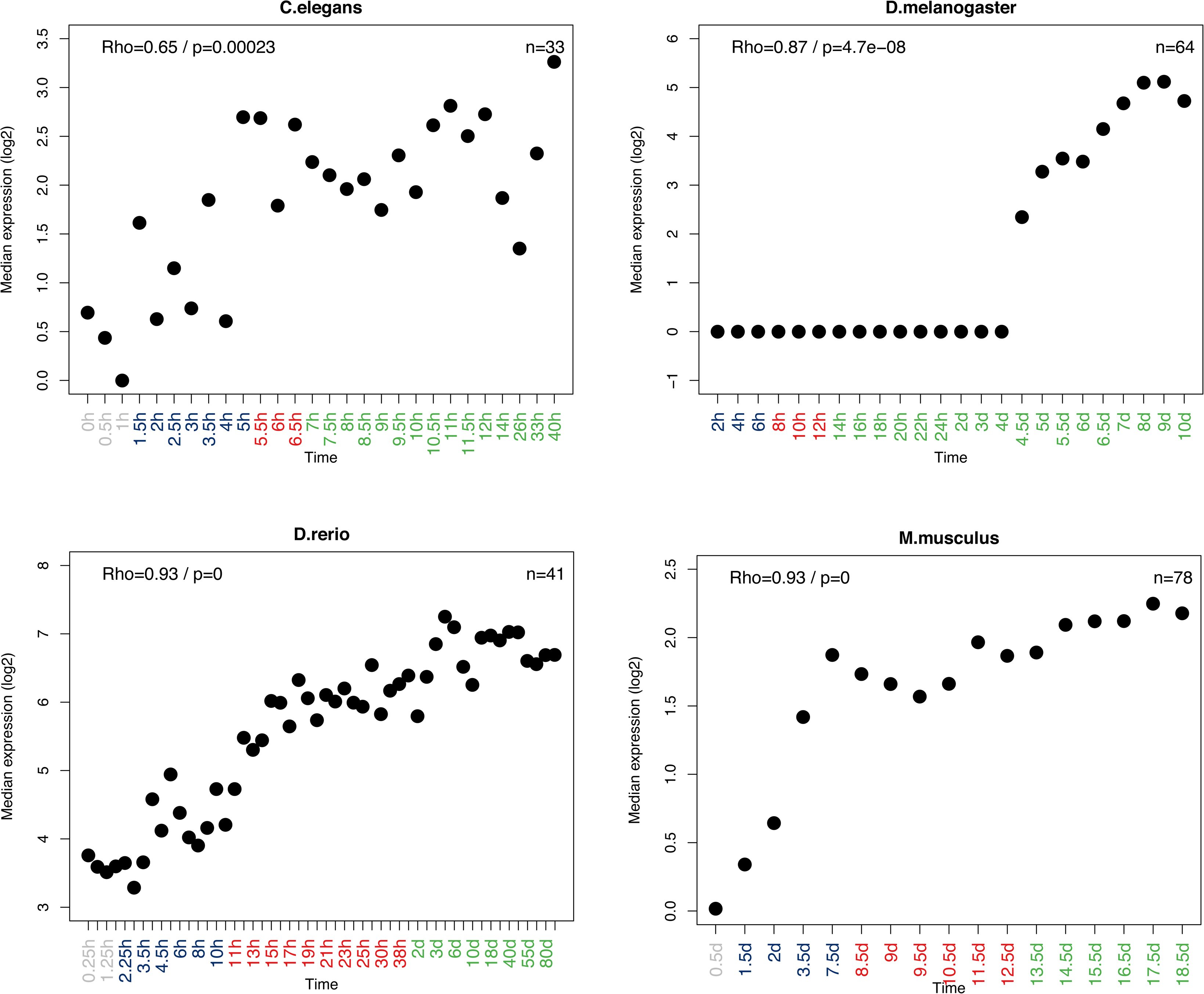
Expression of retrogenes in development. Grey, dark blue, red, and green marked time points in the x-axis represent stages before the start of MZT, early developmental stages, middle developmental stages and late developmental stages respectively. A Spearman correlation was computed between development and median expression. The correlation coefficient (Rho) and *p*-value are indicated in the top-left corner of each plot. The numbers of genes analyzed are noted in the top-right corner of each plot.

These results confirm that late development could allow more transcription of new gene copies, which usually lack efficient regulatory elements and transcriptional activity. Since the first step to functionality is acquiring transcription, we suggest that the functional acquisition and survival at the beginning of life history for new genes could be promoted by expression in late development. When beneficial mutations come, a subset of these new gene candidates could subsequently obtain adaptive functions in late development, evolve efficient regulatory elements, and finally be retained long term in the genome. Thus, the higher proportion of young genes expressed in later development stages can be in part explained by these stages favoring the fixation of new genes.

### Connectivity and dosage imbalance

It has previously been found that, in both *S. cerevisiae* and *C. elegans*, gene duplicability is negatively correlated with protein connectivity (Hughes and Friedman 2005; Prachumwat and Li 2006) which might be explained by dosage balance (Veitia 2002; Papp et al. 2003). Firstly, we checked the relationship of connectivity and duplicability in our datasets. We found, indeed, a negative relationship in *C. elegans* (Figure S11). In *D. melanogaster* and in *D. rerio*, there is a non-monotonous pattern (increasing first, and then decreasing), but the overall trend is more connectivity with less duplicability. In *M. musculus*, however, we did not observe a significant relationship between connectivity and duplicability. Secondly, we calculated a transcriptome index of connectivity (Transcriptome Connectivity Index: TCI). In *C. elegans* and *M. musculus*, earlier developmental stages have higher TCI, which means that these stages trend to have higher expression of more connected genes (Figure 5). In *D. melanogaster* and in *D. rerio*, there is no clear pattern based on individual stages, but the mean TCI of each developmental period (early, middle and late development) also gradually decreases.

**Figure 5:**
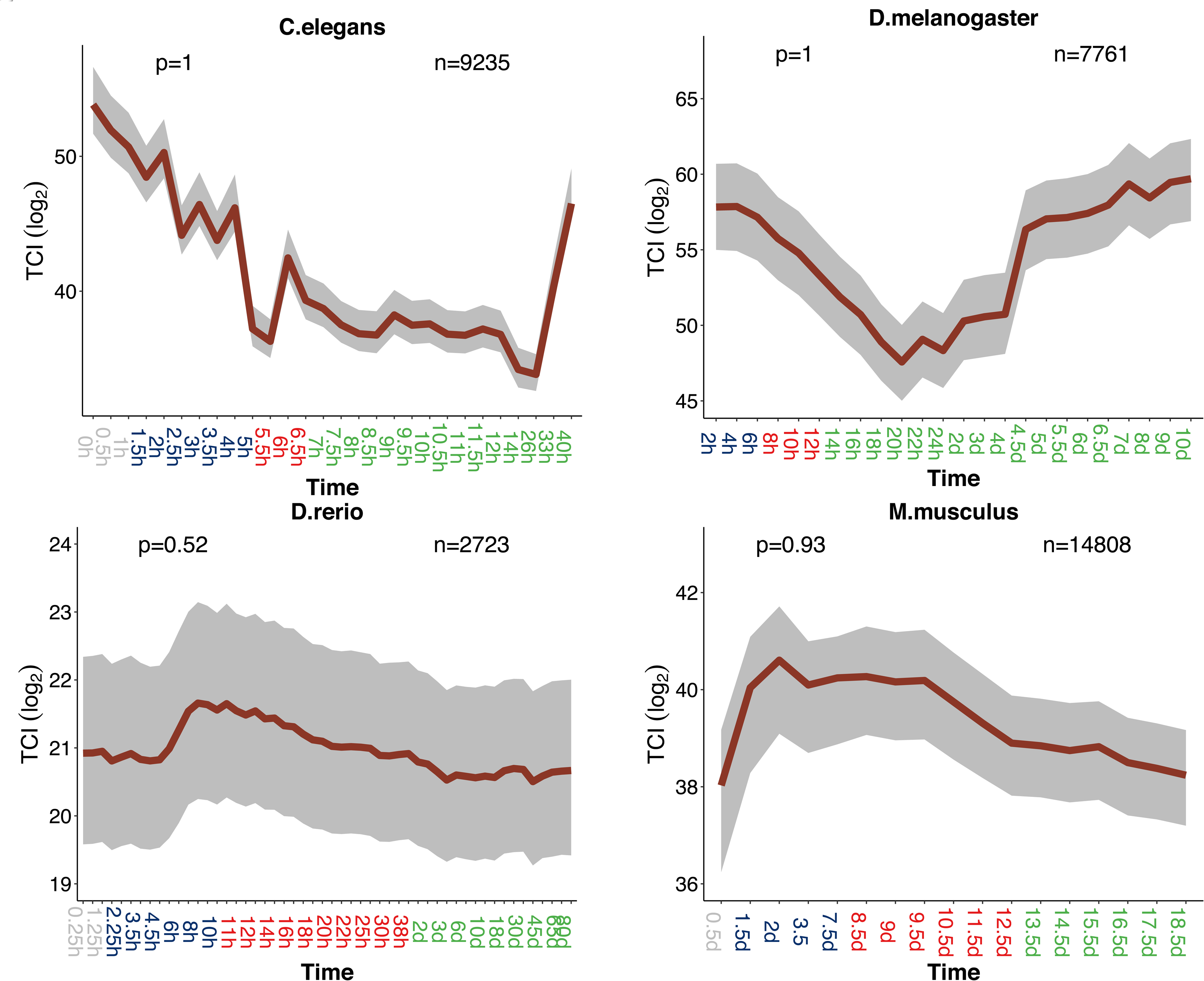
Transcriptome index of connectivity (TCI) across development. Grey, dark blue, red, and green marked time points in the x-axis represent stages before the start of MZT, early developmental stages, middle developmental stages and late developmental stages respectively. TCI is plotted in dark red line. The grey area indicates 95% confidence interval estimated from bootstrap analysis. The *p-*values for supporting the hourglass model (permutation test, early *vs*. middle development) are indicated in the top-left corner of each graph. The numbers of genes analyzed are noted in the top-right corner of each plot.

These results indicate that, at least in *C. elegans*, earlier stages trend to express higher connectivity genes, which are less duplicable because more sensitive to dosage imbalance, but that this cannot be generalized to other animals. Of course, this is not exclusive with an adaptive scenario that early stages lack opportunities for neo- or sub-functionalization, because of simpler anatomical structures, which could also diminish fixation of duplicates in early development.

## Conclusion

Our results concern both patterns and processes of evolution over development. For patterns, we tested the early conservation and hourglass models by using three evolutionary properties: strength of purifying selection, phyletic age and duplicability. The strength of purifying selection on protein sequence supports the hourglass model. Genes under stronger purifying selection are more expressed at middle development stages. Both duplicability and phyletic age support the early conservation model. Less duplicated genes and phyletically older genes are more expressed at earlier stages.

For processes, we investigated the potential causes of the observed patterns. The hourglass pattern of sequence evolution appears to be driven by temporal pleiotropy of gene expression. Genes expressed at middle development evolve under stronger temporal pleiotropic constraints. The enrichment in young phyletic age genes in late development might be related to a testis-like role of late development that facilitates the expression of retrogenes. Finally, in *C. elegans*, connectivity appears to be the main force explaining higher duplicability of genes expressed in later development.

## Materials and Methods

Data files and analysis scripts are available on our GitHub repository: https://github.com/ljljolinq1010/developmental_constraints_genome_evolution

### Expression data sets

#### Main datasets

For *C. elegans* and *D. melanogaster*, we downloaded processed (non-transformed but normalized) RNA-seq data from http://www.stat.ucla.edu/∼jingyi.li/software-and-data.html (Li et al. 2014), which originally comes from (Gerstein et al. 2010; Graveley et al. 2011).

For *D. rerio*, we used the processed (log-transformed and normalized) microarray data from our previous study (Piasecka et al. 2013). This data originally comes from Domazet-Loso and Tautz (2010).

For *M. musculus*, we obtained processed (non-transformed but normalized) RNA-seq data from Hu et al. (2017).

#### Supplementary datasets

For *C. elegans, D. melanogaster* and *D. rerio*, we downloaded the processed (non-transformed and non-normalized) RNA-seq data from Levin et al. (2016). This dataset was generated by CEL-Seq (Hashimshony et al. 2012), a technique for multiplexed single cell RNA-seq. Because CEL-Seq retains only the 3´ end of the transcript, we performed sample normalization based on transcripts per million, but without transcript length normalization. Since only one embryo was sequenced in each stage, there could be larger technical error among lowly expressed genes, like in single cell RNA-seq, especially in early development. So, after normalization, we removed genes with mean expression across samples <1.

For *M. musculus*, the processed (log-transformed and normalized) microarray data was retrieved from Bgee (release 13.1, July 2015; Bastian et al., 2008), a database for gene expression evolution. This data originally comes from (Irie and Kuratani 2011). The detail information of the two datasets listed in Table S1.

#### Omega0 (ω^0^)

The ω_0_ values were downloaded from Selectome (Moretti et al. 2014), a database of positive selection based on the branch-site model (Zhang et al. 2005). Selectome excludes ambiguously aligned regions before model fitting, using Guidance bootstrapping and M-Coffee consistency. Omega0 is the dN/dS ratio (dN is the rate of non-synonymous substitutions, dS is the rate of synonymous substitutions) of the subset of codons which have evolved under purifying selection according to the branch-site model. We used ω_0_ from the Clupeocephala branch, the Murinae branch, and the Melanogaster group branch for *D. rerio, M. musculus*, and *D. melanogaster*, respectively. One gene could have two ω_0_ values in the focal branch because of duplication events. In this case, we keep the value of the branch following the duplication and exclude the value of the branch preceding the duplication.

#### Phyletic age data

Phyletic ages were retrieved from Ensembl version 84 (Yates et al. 2016) using the Perl API. For each gene, we browsed its gene tree from the root and dated it by the first appearance. We assigned the oldest genes with phyletic age value of 1 and the youngest genes with the highest phyletic age value. So, genes can be split into discrete “phylostrata” by phyletic age. We classified 3 phylostrata, 4 phylostrata, 9 phylostrata and 18 phylostrata respectively for *C. elegans, D. melanogaster, D. rerio* and *M. musculus*. The definition of phylostrata is dependent on the available genome sequences in related lineages, hence the differences between species.

#### Number of paralogs

We retrieved the number of paralogs from Ensembl release 84 (Yates et al. 2016) using BioMart (Kinsella et al. 2011).

#### Retrogene data

For *C. elegans,* we retrieved 33 retrogenes from Zou et al. (2012). For *D. melanogaster,* we retrieved 72 retrogenes from retrogeneDB (Kabza et al. 2014). For *D. rerio* we retrieved 113 retrogenes from Fu et al. (2010). For *M. musculus* we retrieved 134 retrogenes from Potrzebowski et al. (2008).

#### Connectivity data

We retrieved connectivity (protein-protein interactions) data from the OGEE database (Chen et al. 2012).

#### Testis specific genes

We first retrieved processed (normalized and log-transformed) RNA-seq data of 22 *M. musculus* tissues and 6 *D. melanogaster* tissues from Kryuchkova-Mostacci and Robinson-Rechavi (2016a).

Then, we calculated tissue specificity based on Tau (Yanai et al. 2005; Kryuchkova-Mostacci and Robinson-Rechavi 2016b):

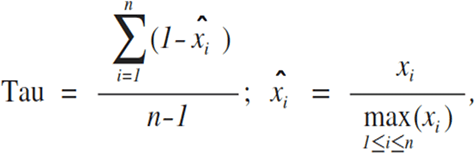

where *n* is the number of tissues, and *x*_*i*_ is the expression of the gene in tissue *i*. This index ranges from zero (broadly expressed genes) to one (genes specific to one tissue). All genes that were not expressed in at least one tissue were removed from the analysis.

Finally, we defined genes with highest expression in testis and with tissue specificity value ≥ 0.8 as testis specific genes.

#### Transcriptome index analysis for different evolutionary parameters

The TEI (transcriptome evolutionary index) is calculated as:

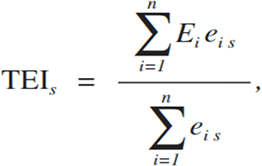

where *s* is the developmental stage, *E*_*i*_ is the relevant evolutionary parameter (ω_0_, paralog number, phyletic age, stage specificity, or protein connectivity) of gene *i, n* is the total number of genes, and *e*_*is*_ is the expression level of gene *i* in developmental stage *s*; by default we use log-transformed expression levels for *e*_*is*_.

#### Confidence interval analysis

Firstly, we randomly sampled gene IDs from each original data set 10,000 times with replacement. Then, we computed transcriptome indexes for the 10,000 samples. Finally, the 95% confidence interval is defined as the range from quantile 2.5% to quantile 97.5% of the 10,000 transcriptome indexes. This approach was integrated into myTAI (Drost et al. 2017), a R package for evolutionary transcriptome index analysis.

#### Stages before the start of maternal to zygote transition

The stages before the start of the maternal to zygote transition (MZT) were defined from Tadros and Lipshitz (2009). For C. elegans and *D. melanogaster*, it’s the first 1 hour of embryo development; for *D. rerio*, it’s the first 2 hours of embryo development; for *M. musculus*, it’s the first day of embryo development.

#### Phylotypic period

From both morphological and genomic studies, we defined the phylotypic period of each species as follows: for *C. elegans*, the phylotypic period is defined as the ventral enclosure stage (Levin et al. 2012); for *D. melanogaster*, the phylotypic period is defined as an extended germband stage (Sander 1983; Kalinka et al. 2010); for *D. rerio*, the phylotypic period is defined as the segmentation and pharyngula stages (Ballard 1981; Wolpert 1991; Slack et al. 1993; Domazet-Loso and Tautz 2010); for *M. musculus*, the phylotypic period is defined as Theiler stages 13 to 20 (Ballard 1981; Wolpert 1991; Slack et al. 1993; Irie and Kuratani 2011).

#### Permutation test

We first assigned all development stages to three broad development periods (early: after the start of MZT and before the phylotypic period, middle: the phylotypic period, and late: after the phylotypic period). Next, we calculated the difference of mean transcriptome indexes between the early module and the middle module (Δ*e-m*). Then, we permuted the values of the relevant parameter (ω_0,_ paralog number, phyletic age, stage specificity or protein connectivity) 10,000 times. Finally, we approximated a normal distribution for Δ*e-m* based on 10,000 Δ*e-m* values computed from the permutated samples. The *p-*value of the hourglass model *vs*. the early conservation model for each parameter is the probability of a randomly sampled Δ*e-m* exceeding the observed *Δ*e*-m*. For protein connectivity, the *p-*value of the hourglass model is the probability that a randomly sampled Δ*e-m* lower than the observed *Δe-m*.

## Acknowledgements

We thank Marie Zufferey and Nadezda Kryuchkova for help with preliminary work. We thank the Bgee team, Marie Zufferey, Nadezda Kryuchkova and Barbara Piasecka for help with data retrieval, Julien Roux for help with the Ensembl API, Kamil Jaron and Andrea Komljenovic for help with programing, all members of the Robinson-Rechavi lab and Hajk-Georg Drost for helpful discussions. Part of the computations were performed at the Vital-IT (http://www.vital-it.ch) centre for high-performance computing of the SIB Swiss Institute of Bioinformatics. This work was supported by Swiss National Science Foundation grant 31003A_153341 / 1.

## Author contributions

JL and MRR designed the work. JL performed the data gathering and analysis. JL and MRR interpreted the results. JL wrote the first draft of the paper. JL and MRR finalized the paper.

## Figures legends

**Figure S1:**
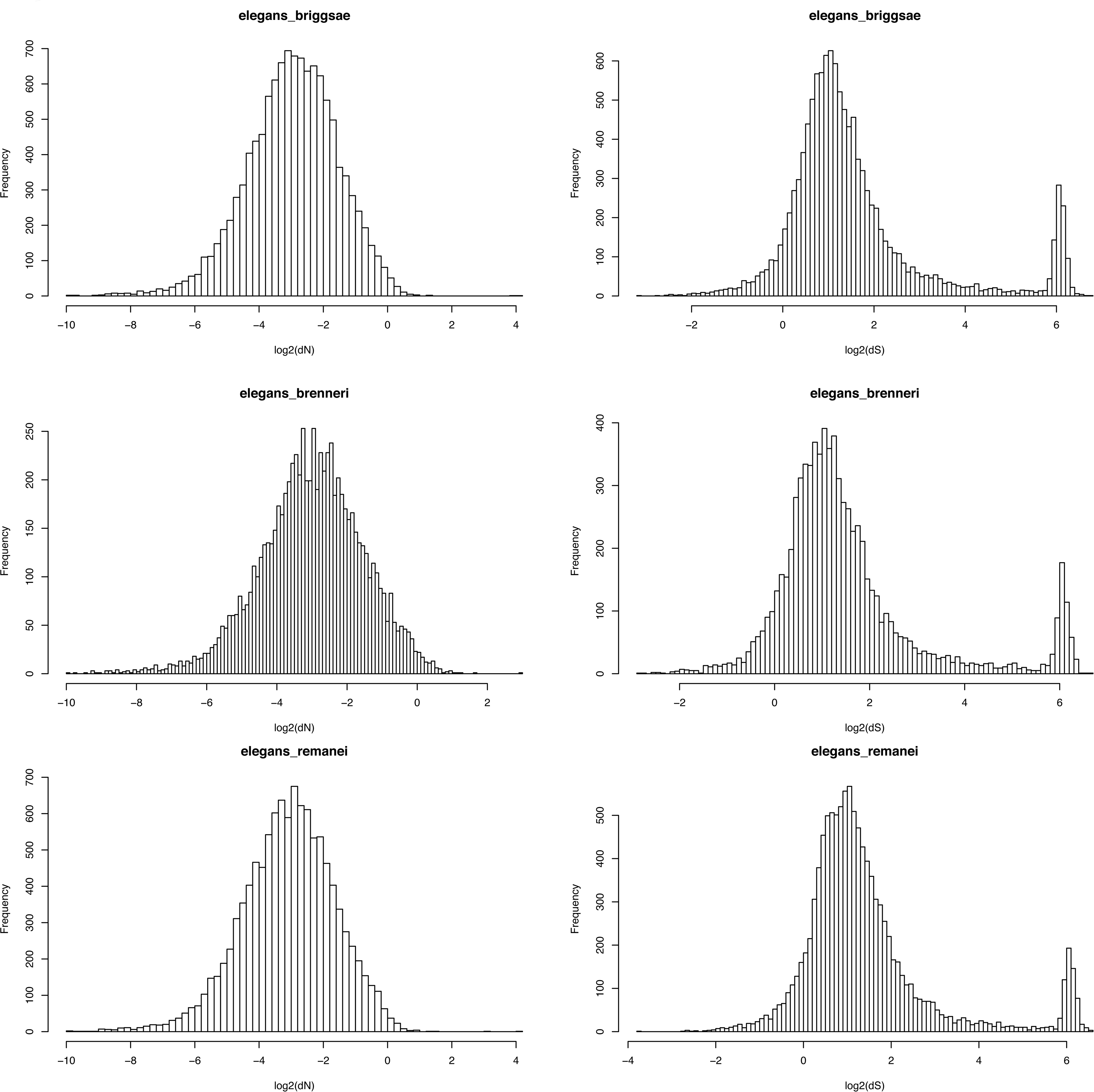
Distributions of dN and dS for *C. elegans.*

**Figure S2:**
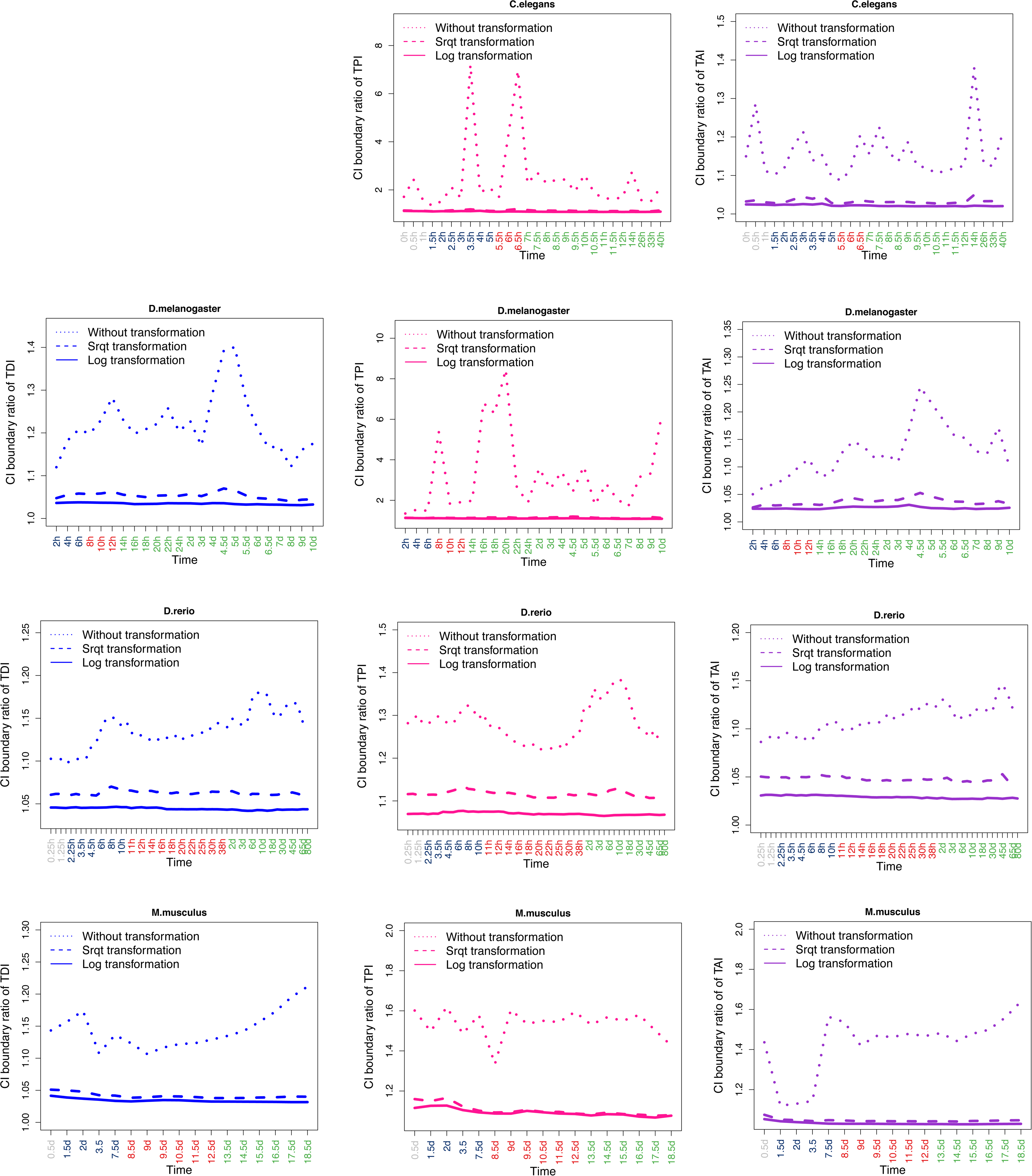
Comparison of 95% confidence intervals from transformed and non-transformed expression values. Grey, dark blue, red, and green marked time points in the x-axis represent stages before the start of MZT, early developmental stages, middle developmental stages and late developmental stages respectively. Y-axis represents the ratio of upper to lower 95% confidence interval boundary. The ratio from non-transformed expression values is plotted in dotted lines, while the ratio from log_2_ transformed expression values is plotted in solid lines, and the ratio from square root (abbreviated as “sqrt”) transformed expression values is plotted in dashed lines.

**Figure S3:**
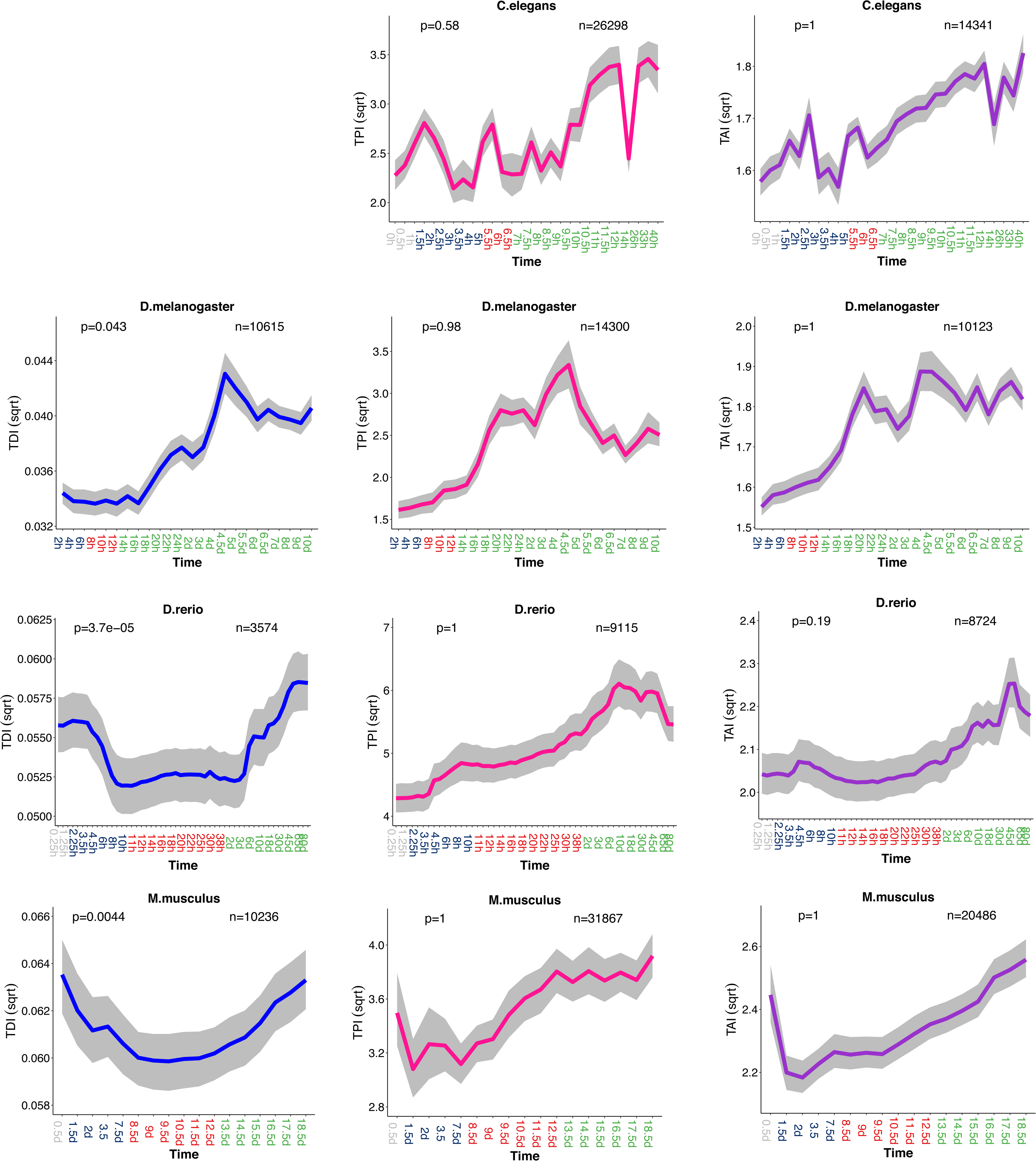
Evolutionary transcriptome indexes based on square root transformed expression values. Legend as Figure 1, but here the indexes are based on square root transformed expression values.

**Figure S4:**
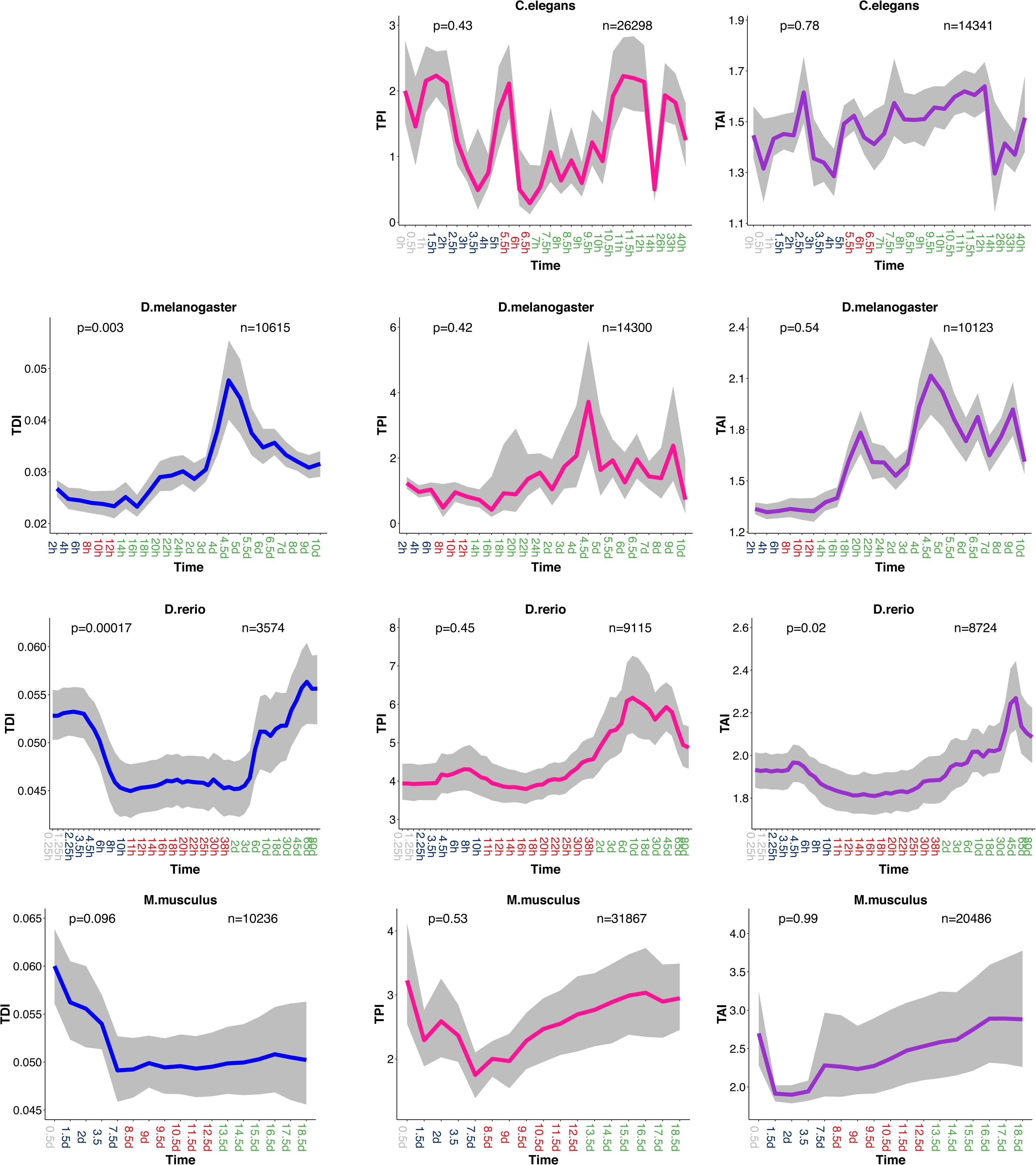
Evolutionary transcriptome indexes based on non-transformed expression values. Legend as Figure 1, but here the indexes are based on non-transformed expression values.

**Figure S5:**
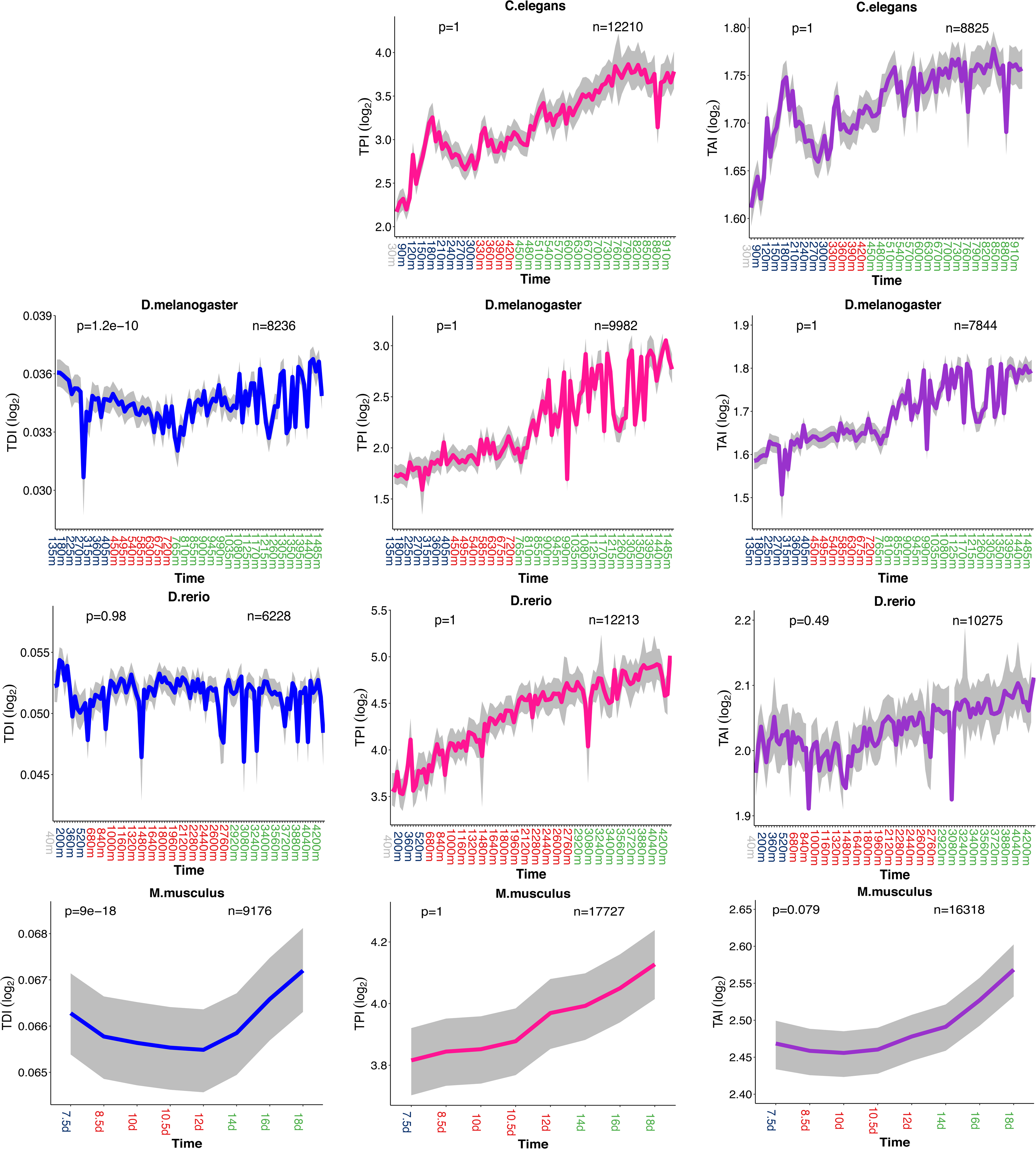
Evolutionary transcriptome indexes based on supplementary datasets. Legend as Figure 1, but here the indexes are based on supplementary datasets.

**Figure S6:**
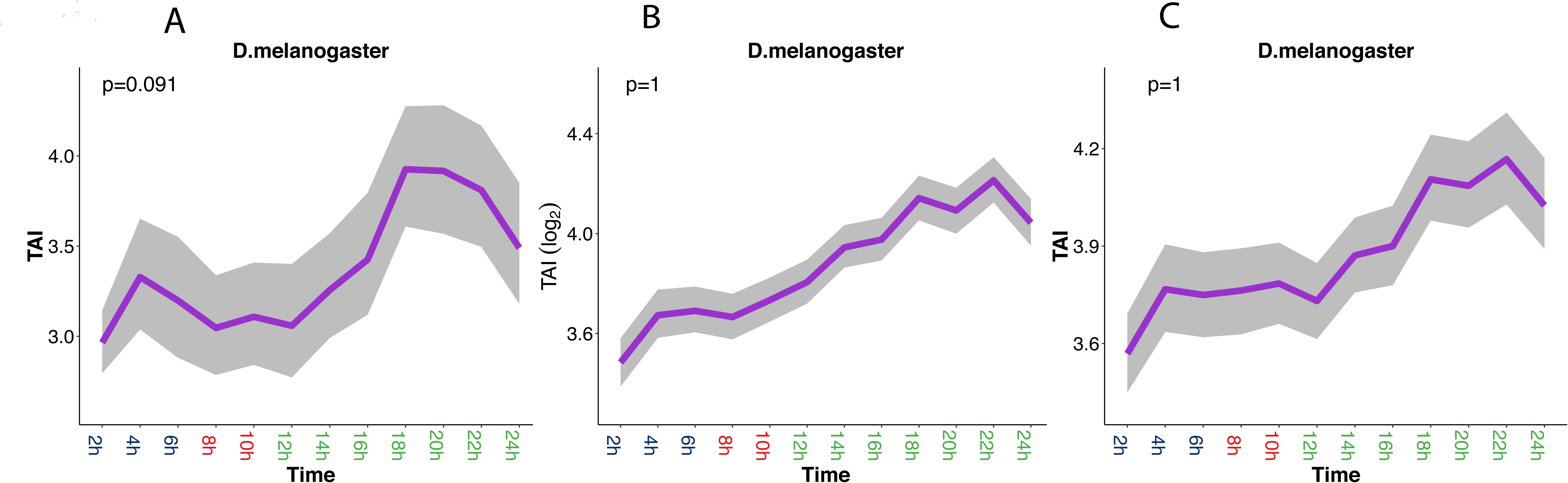
Comparison of transcriptome phyletic age indexes (TAI). Dark blue, red, and green marked time points in the x-axis represent early developmental stages, middle developmental stages and late developmental stages respectively. TAI is plotted in purple line. The grey area indicates 95% confidence interval estimated from bootstrap analysis. The *p-*values for supporting the hourglass model (permutation test, early *vs*. middle development) are indicated in the top-left corner of each graph. A: TAI based on non-transformed expression values. B: TAI based on log_2_ transformed expression values. C: TAI based on non-transformed expression values, excluding the top 10% highest expressed genes.

**Figure S7:**
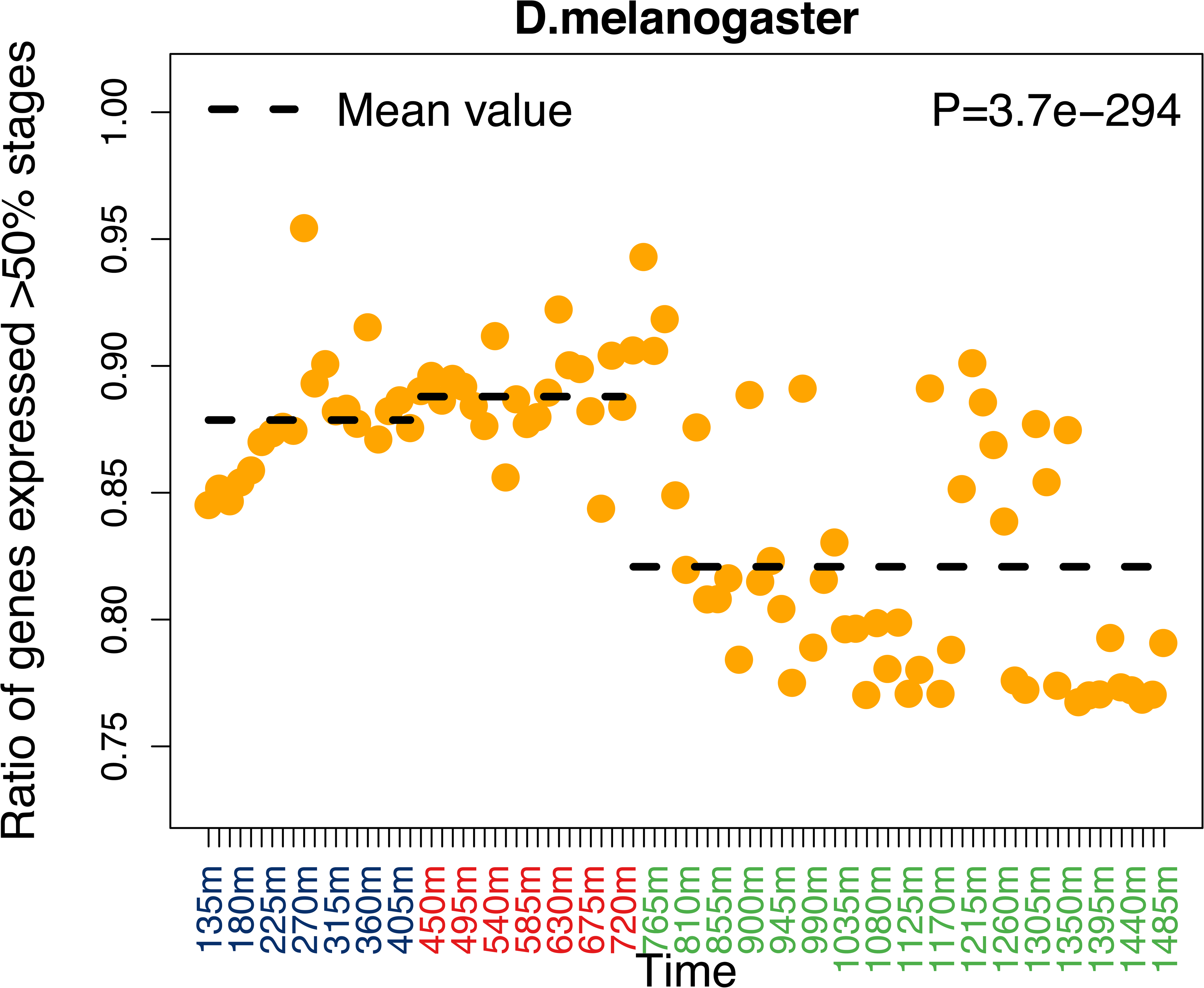
Proportion of temporal pleiotropic genes for *D. melanogaster* with the supplementary dataset. Legend as Figure 2, but here the result comes from the supplementary dataset of *D. melanogaster*.

**Figure S8:**
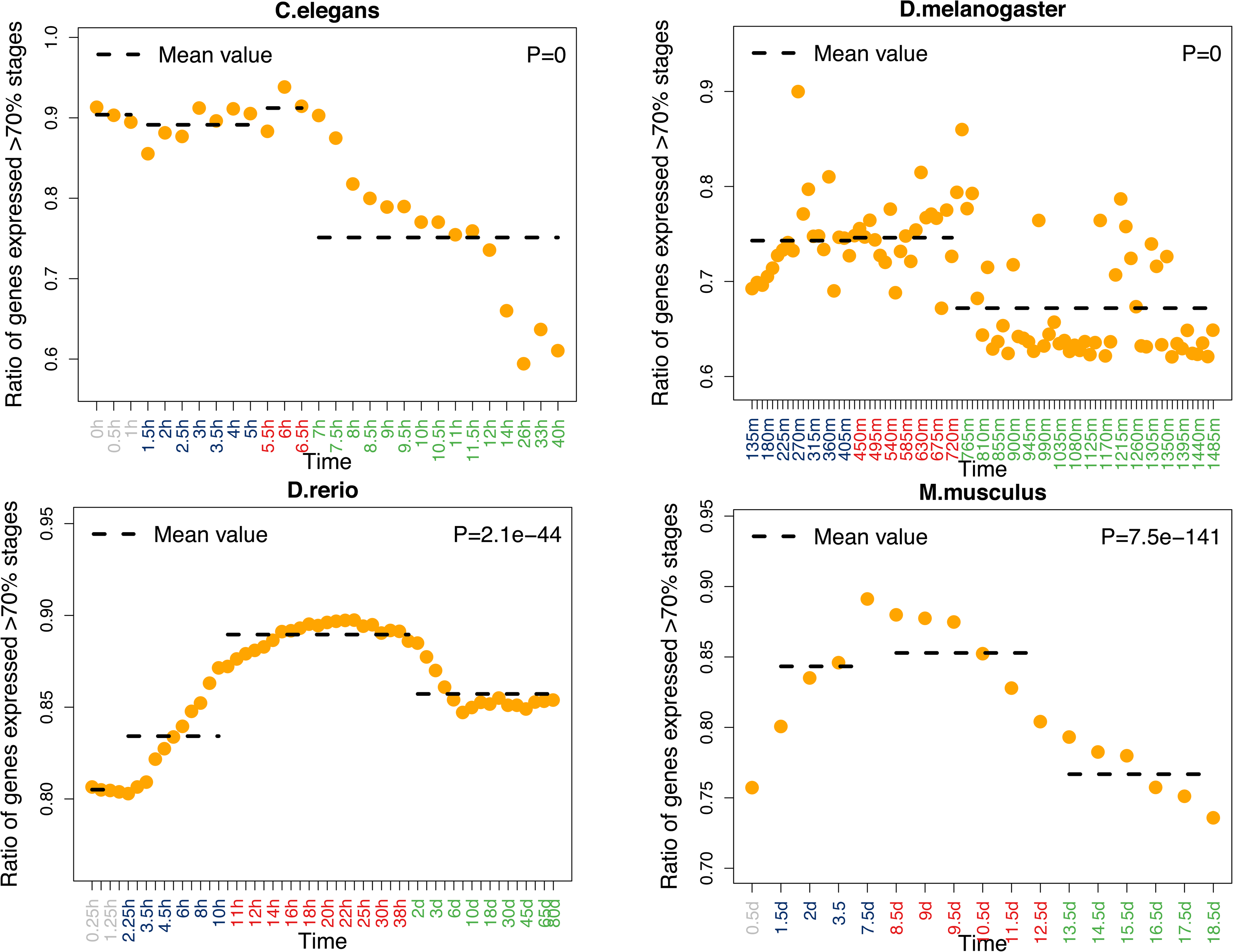
Proportion of temporal pleiotropic genes defined as expressed in more than 70% stages. Legend as Figure 2, but here the temporal pleiotropic genes are defined as expressed in more than 70% of stages.

**Figure S9:**
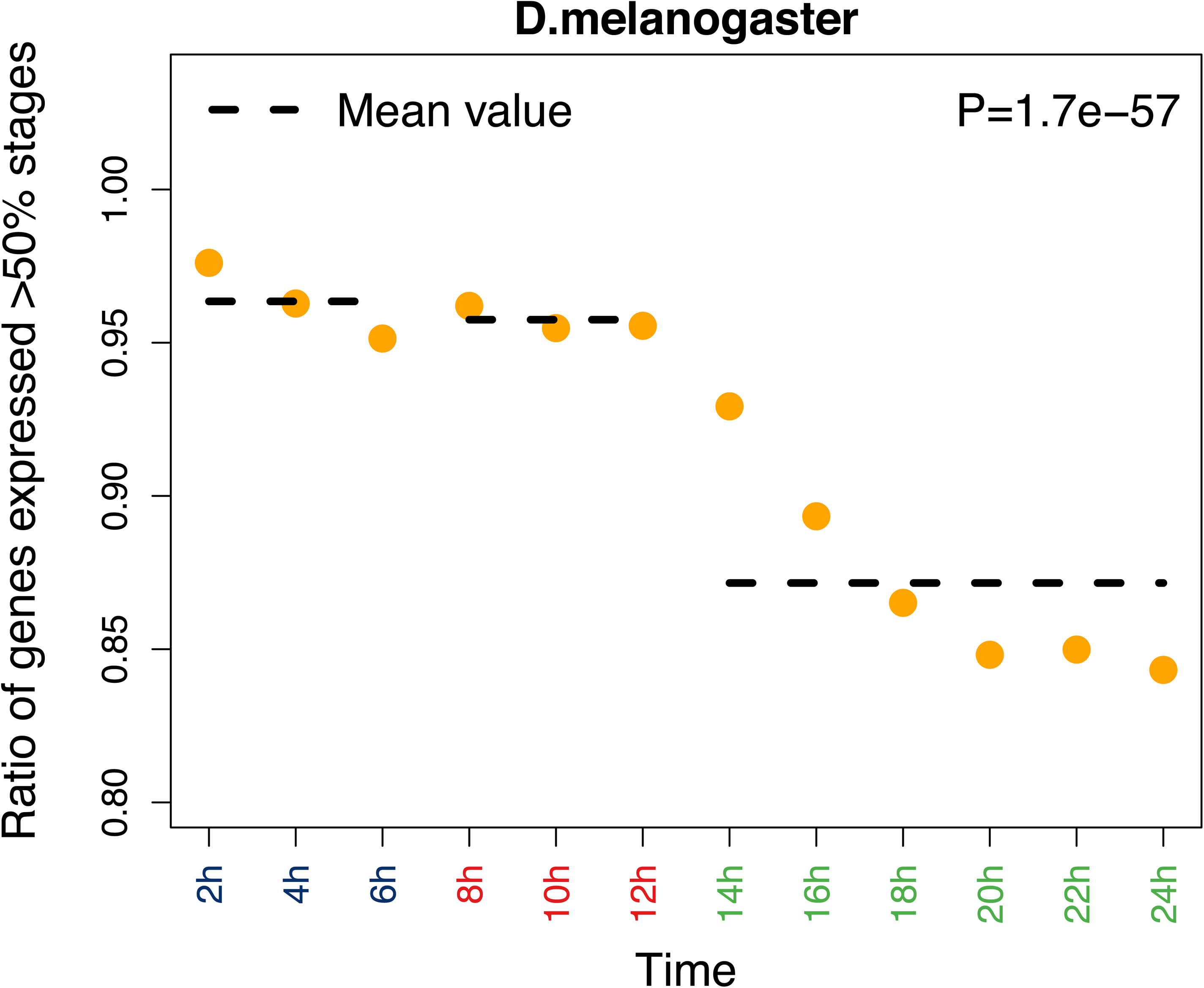
Proportion of temporal pleiotropic genes for *D. melanogaster* after removal of the second period of late development. Legend as Figure 2.

**Figure S10:**
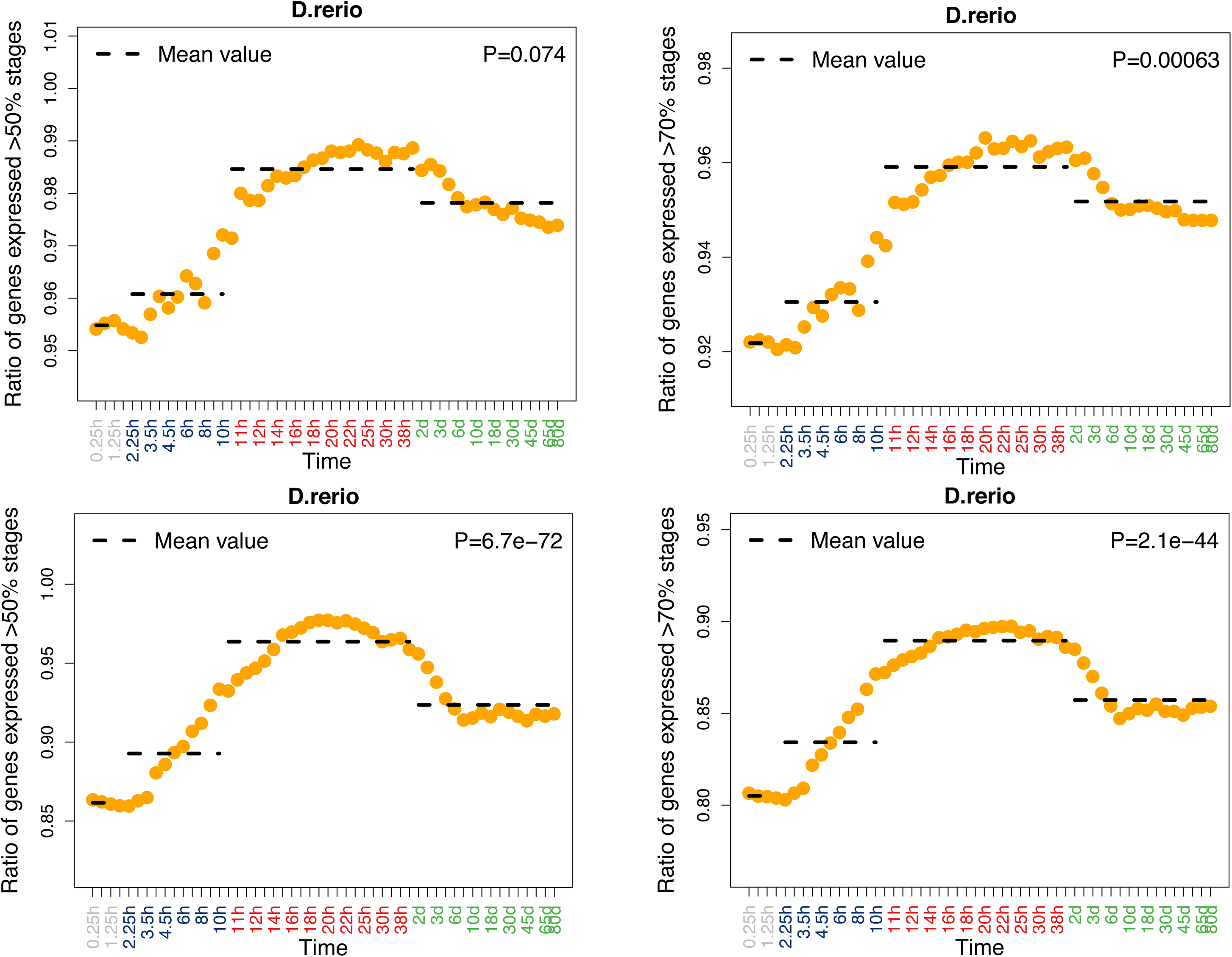
Proportion of temporal pleiotropic genes for *D. rerio* based on expressed genes defined as microarray signal rank in top 90% or 50%. Legend as Figure 2. In upper panel graphs, expressed genes defined as microarray signal rank in top 90%. In lower panel graphs, expressed genes defined as microarray signal rank in top 50%.

**Figure S11:**
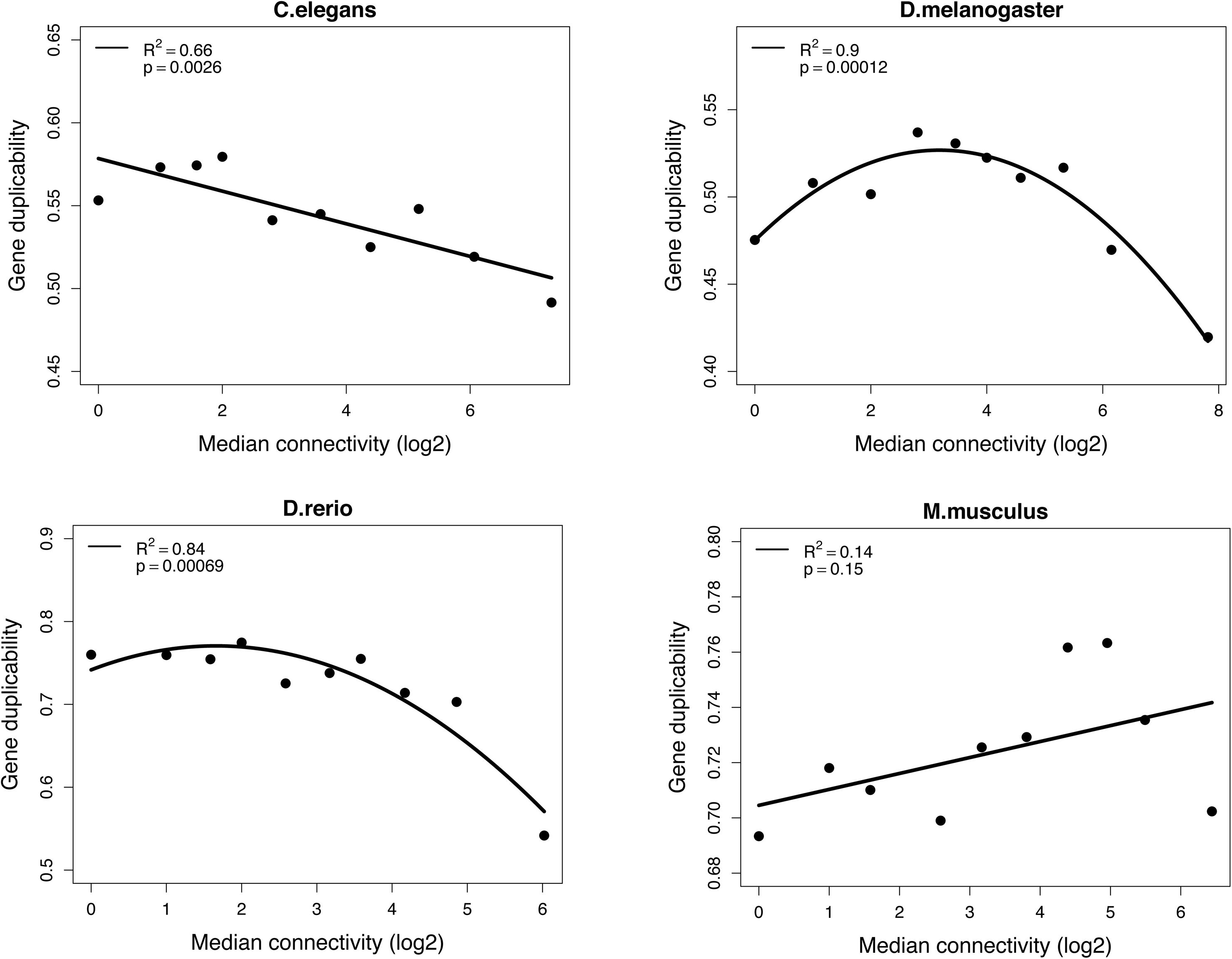
Relation of protein connectivity and duplicability. Genes were split into 10 bins according to their connectivity. The duplicability in each bin was measured by the number of genes with paralogs divided by the number of all genes. The duplicability was fit by regression (the first degree of polynomial for *C. elegans* and *M. musculus,* while the second degree of polynomial for *D. melanogaster* and *D. rerio*), whose *R^2^* and *p-*value are indicated in the top-left corner of each graph. The median connectivity of each bin was plotted on the x-axis (in log_2_ scale).

**Table S1.** Expression datasets used in this study; “main”: dataset used for the figures in the manuscript; “supplementary”: dataset used for the figures in the supplementary materials.

| Species | *C. elegans* |  | *D. melanogaster* |  | *D. rerio* |  | *M. musculus* |  |
| --- | --- | --- | --- | --- | --- | --- | --- | --- |
| Dataset | main | supplementary | main | supplementary | main | supplementary | main | supplementary |
| ref | (Gerstein et al. 2010) | (Levin et al. 2016) | (Graveley et al. 2011) | (Levin et al. 2016) | (Domazet-Loso and Tautz 2010) | (Levin et al. 2016) | (Hu et al. 2017) | (Irie and Kuratani 2011) |
| datatype | RNA-seq | RNA-seq | RNA-seq | RNA-seq | Microarray | RNA-seq | RNA-seq | Microarray |
| replicates | no | no | no | no | 2 | no | $\geq 2$ | $\geq 2$ |
| stages before MZT | 3 | 2 | 0 | 0 | 4 | 3 | 1 | 0 |
| early stages | 7 | 23 | 3 | 20 | 11 | 13 | 4 | 1 |
| middle stages | 3 | 12 | 3 | 20 | 21 | 55 | 6 | 4 |
| embryo late stages | 7 | 49 | 6 | 51 | 3 | 35 | 6 | 3 |
| post-embryo late stages | 8 | 0 | 12 | 0 | 13 | 0 | 0 | 0 |
| total stages | 28 | 86 | 24 | 91 | 52 | 106 | 17 | 8 |

